# Capturing Neonatal Cardiomyocyte – Endothelial interactions on-a-Chip

**DOI:** 10.1101/2020.08.10.243600

**Authors:** Hossein Tavassoli, Young Chan Kang, Prunella Rorimpandey, John E Pimanda, Vashe Chandrakanthan, Peggy P.Y. Chan

## Abstract

The neonatal heart has been the focus of numerous investigations due to its inherent regenerative potential. However, the interactions between neonatal cardiomyocytes (CMs) and endothelial cells (ECs) have been difficult to model and study due to the lack of an appropriate device. Here, we developed a method to culture primary neonatal CMs and ECs in a microchip and characterise their behavioural properties over a 14-day period. By implementing cell migration analyses coupled with immunostaining and confocal microscopy, we were able to identify and quantify sub-populations of migratory and non-migratory ECs. In CM–EC co-cultures, migrating ECs were found to move in higher numbers and longer distances compared to migrating CMs. In the presence of CMs, non-migrating ECs established connexin gap junctions and formed CM–EC cell aggregates, which were likely a priming event for endothelial organoid formation. This microfluidic device also enabled us to visualise the temporal sequence organoid formation and phenomena such as collective cell migration, CM–EC trans-differentiation and synchronisation of CM beating. This microchip based culture system has potential applications for tissue engineering and drug discovery.

## 1. Introduction

Heart disease remains the leading cause of mortality and morbidity in the world ^[1]^. Understanding the cellular events that govern heart tissue development could be useful for the design and development of therapeutics to restore cardiac function. Much of our current understanding of cellular events associated with cardiac development has been gleaned from *in vitro* dish ^[2]^, plate ^[3]^ or trans-well based co-culture systems ^[4,5]^ and *in-vivo* animal models ^[6,7]^.

*In vivo* cardiac cell biology studies and disease-based animal models are expensive and laborious. They are also not amenable to tracking individual cell behaviour. For example, cardiomyocytes (CMs) and endothelial cells (ECs) are scattered throughout the endocardium and myocardium ^[8,9]^, and it is challenging to pinpoint the fate of individual cells within a thick complex tissue. This is one of the reasons why the morphogenetic steps involved in CMs and ECs migration and differentiation during cardiac development are still not well defined ^[10–12]^. It is even harder to identify precise cell-cell interactions between CMs and ECs in this setting.

Although more amenable to controlled experimentation, a major drawback of conventional *in vitro* culture systems is that they cannot mimic the *in vivo* cellular microenvironment, as these cell cultures often lose their intercellular contacts ^[13]^ and differentiation functions ^[14]^. In recent years, new *in vitro* models such as scaffold systems ^[15]^, pre-formed vascular networks ^[16]^, cell sheet engineering technologies ^[17]^, and micro engineered systems ^[18]^ have been developed with the goal of mimicking the complex architecture of *in vivo* models. Amongst these various *in vitro* culture systems, microfluidic devices have shown promise as *in vitro* miniature models of native tissues ^[19]^. As the regenerative capacity of the murine heart decreases after postnatal day 7 ^[20]^, much of our current knowledge pertaining to cardiac tissue development has come from studying mouse embryonic and neonatal hearts ^[20,21]^. CD31 expressing ECs are a major constituent of the coronary vasculature ^[22]^. Several groups have investigated the co-culture of CMs and ECs ^[12,23–25]^. These studies have involved the use of adult CMs or CMs derived from either induced pluripotent stem cells (iPSC) ^[26]^, or human embryonic stem cells (hESC) ^[27]^. Other reports have used ECs derived from extra-cardiac tissues such as umbilical vein ^[28]^, peripheral blood ^[29]^, adult ECs ^[30]^ or ESC-derived ECs ^[28]^. Yet, due to sensitivity and complexities present in isolation and culture of primary cardiac cells, there are no reports of using a platform to investigate the crosstalk between primary beating neonatal CMs and CD31+ cardiac ECs from neonates– a developmental time point at which CMs retain their regenerative potential.

In this study, we first established a method to isolate and culture neonatal beating CMs and CD31+ cardiac ECs on a chip, then used this chip to study the behaviour of neonatal CMs and neonatal ECs, and demonstrate cell-cell interactions between CMs and ECs.

## 2. Results and Discussion

### 2.1. Neonatal heart-on-a-chip: overview of experimental design

In order to gain insights into CM and EC behaviour during neonatal cardiac development, a microfluidic device containing three parallel channels was built. A detailed workflow of the set up and quantification analysis is shown in **Figure 1**. Wild type C57BL/6 mice were time mated to generate wild-type neonates (**Figure 1 A**). Whole hearts of two day old neonates (P2) were dissected and dissociated to generate cell suspensions that contained a heterogeneous cardiac cell population. Methods were developed to further isolate CMs and ECs from this heterogeneous pool and subsequently culture them in the microfluidic device (**Supporting information**, **Figure S1-S3** and **Video S1-S2**). CMs and ECs were separately seeded into one of the side channels precoated with matrigel. The middle channel was filled with matrigel to mimic the 3-dimensional extracellular matrix (ECM) environment in neonatal heart tissue (**Figure 1 B**). Cells from each of the side channels were allowed to grow and migrate into the middle channel over a period of 7-14 days. Cells cultured in each of the side channels could receive soluble factors secreted by cells from the opposite side channel through the middle ECM channel. Imaging and cell counting analyses were performed to monitor cells in the ECM at day 0, 7, and 14 (**Figure 1 C**). The migratory behaviour of cells was quantified by measuring the following three parameters: number of migrating cells, migration distance (measured as Euclidean distance) and migration rate (**Figure 1 D).** Three regions of interest (ROI) were randomly selected from each chip (**Figure 1 E**). Cells that moved into the middle channel from any of the side channels were defined as migrating cells. Cells that stayed within any of the side channels were defined as non-migrating cells.

**Figure 1.**
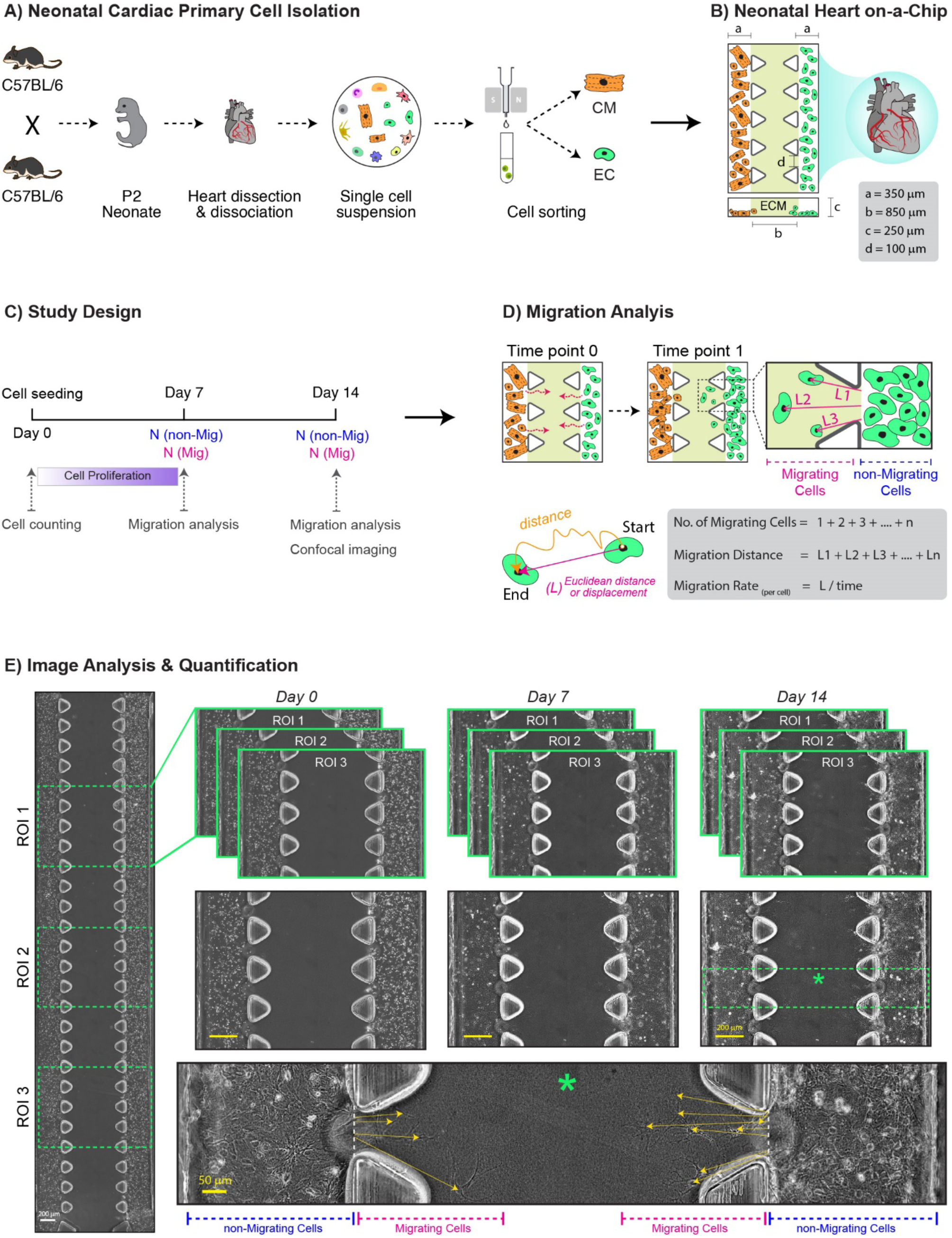
Overview of microchip design and cell migration analysis workflow. A) Primary neonatal (P2) cardiac cells were isolated and sorted into distinct populations of cardiomyocyte and endothelial cells using MACS. **B)** Schematic illustration showing the design and dimensions of the microfluidic device. Cells were loaded in each of the side microchannels. The middle channel was filled with ECM mimetic matrigel that allows cells to migrate in a 3D environment. The width of the side channels was 350 micrometres and that of the middle channel 850 micrometres. The distance between each post was 100 micrometres and the height of each channel was 250 micrometres. **C)** The timeline and design of the study. Number of migrating and non-migrating cells were counted after 7 and 14 days of culture (N(Mig) and N(non-Mig)). Quantitative migration analysis and confocal microscopy was performed after 7 and 14 days of culture. **D)** Schematic and listed formulas showing quantification of cell migration from side channels into the middle channel at each time point. **E)** Representative tile-scan image of a co-culture microchip (configuration e) after cell seeding at day 0. Three regions of interest (ROI) were selected from each microfluidic chip in an unbiased way. The magnified area of a selected ROI at day 0 shows even distribution of cells in the side channels. Whereas at the end of 7 and 14 days, some cells migrated from the side channels into the middle channels, these cells were denoted as migrating cell. The majority of cells remained in the side channels and were denoted as non-migrating cells. Image J software was used for the image analysis; the interface between side channel and middle channel was chosen as the starting point for cell migration, the location of each cell in the middle channel was determined as the end point for cell migration. The Euclidean distance was determined as the shortest distance between the start and end point for a specific cell.

### 2.2. Migration of neonatal CMs and ECs on a microchip

In all experiments, neonatal CMs and ECs were isolated from P2 heart tissues, cultured in microchips and analysed (**Figure 2 A**). To investigate whether the neonatal cells migrate randomly into the hydrogel or migrate via cell-to-cell signalling process, a set of control experiments (**Figure 2 B (a-d)**) were devised to test the migration tendency of neonatal cells in mono-culture, where cells of a particular type were injected into one or both side channels. In a separate set of experiments, the impact of mixed CM/EC cultures on the migration of CMs and ECs was also assessed. To this end, three co-culture configurations were used: (i) CMs were cultured in the left side channel while ECs were cultured in the right side channel (**Figure 2 B (e)),** (ii) CMs were cultured in the left side channel while a mixture of CMs and ECs were co-cultured in the right side channel (**Figure 2 B (f)**), (iii) ECs were cultured in the left side channel while a mixture of CMs and ECs were co-cultured in the right side channel (**Figure 2 B (g)**).

**Figure 2.**
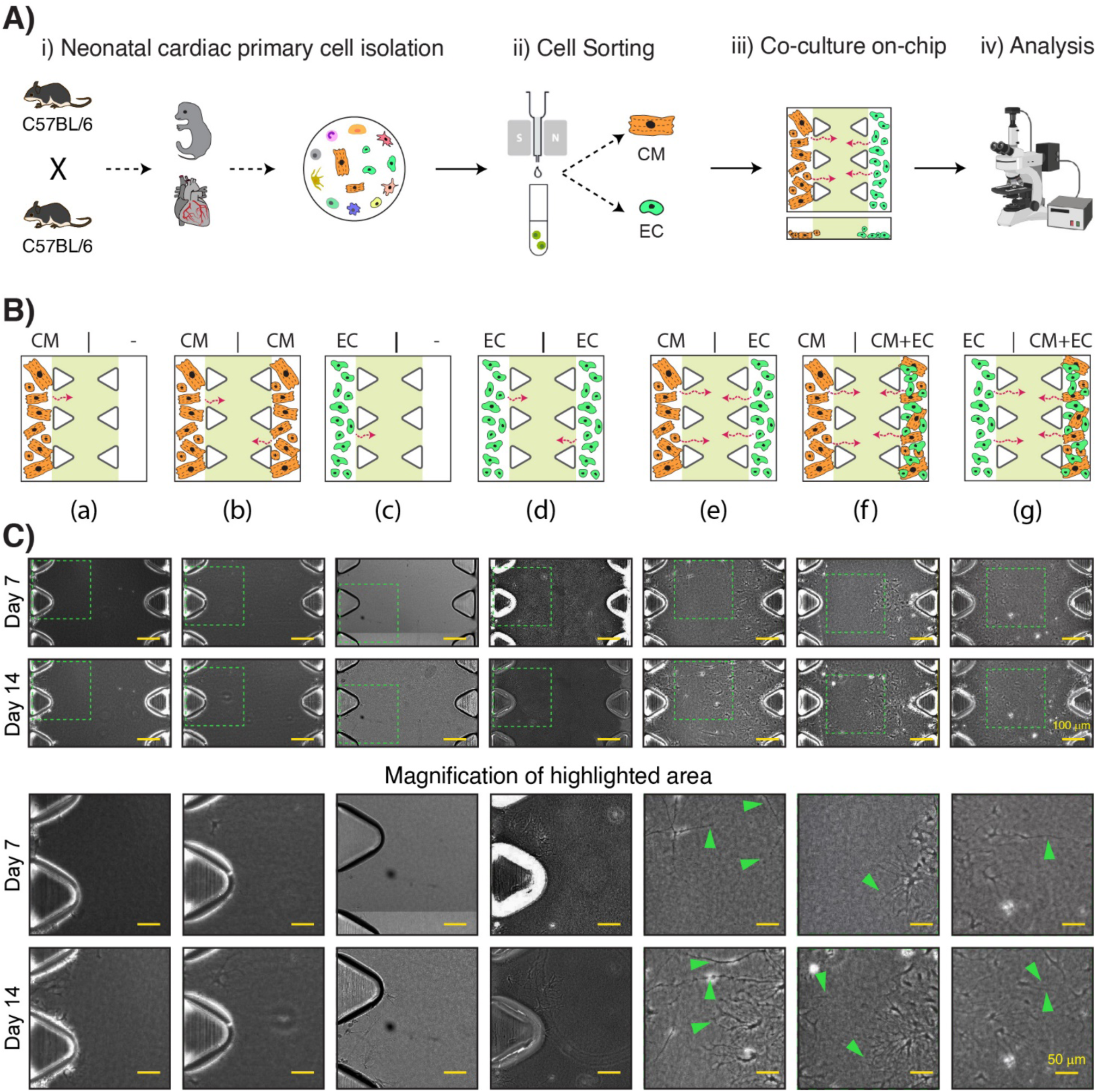
Morphology of neonatal cardiomyocyte and endothelial cells in co-culture microfluidic devices. **A)** Schematic representation of the experimental design: i) Primary neonatal (P2) cardiac cells were isolated and ii) sorted into distinct populations of cardiomyocyte and endothelial cells via MACS. iii) Specific cell populations were loaded into a microfluidic device and iv) Migration analysis was performed using phase contrast and confocal microscopy. **B)** Schematic representation showing microchips with different configurations. (a – d) Monoculture: control experiments that test the migration tendency of a population that contains only one cell type, (e – f) Coculture: experiments that test the migration tendency of populations that contain two types of cells in different configurations. **C)** Representative phase contrast images showing migration of primary neonatal cardiomyocyte and endothelial cells into the middle ECM microchannel after 7 and 14 days of culture. Dotted line highlighted randomly selected ROI. The selected ROIs were shown in higher magnification at the two bottom image panels. Green arrows point to the tunnelling nanotubes (TNT) like structures found in-between cells.

When CMs were cultured in only one of the side channels in monoculture, very few CMs migrated (**Figure 2 C a**). Similar behaviour was observed when both side channels were cultured with CMs (**Figure 2 C b**). Unlike CMs, small numbers of ECs were found to migrate into the middle channel when ECs were cultured as monoculture (**Figure 2 C (c-d)**) regardless of whether ECs were cultured on one or both of the side channels. With co-cultures, as expected, some ECs migrated into the ECM channel. However, compared to monocultures, there were more migrating ECs in co-cultures regardless of the co-culture configuration (**Figure 2 C (e-g)**). In contrast to CMs in monocultures, more CMs were found to migrate into the ECM channel in the co-culture setting. Microscopic images revealed that some of the migrating cells had formed tunnelling nanotube (TNT) like structures in the ECM channel (**Figure 2 C (e-g;** green arrowheads). TNT formation between ECs and CMs are known to occur as a conduit for long-range communication ^[31–33]^ and indicated that the migrating cells had established communication conduits between themselves.

Interestingly, not all cells from the side channel had migrated into the middle ECM channel. We therefore quantified the number of non-migrating cells and migrating cells in different configurations (**Figure 3 A)**. In monoculture experiments with cells cultured on only one of the side channels (configurations (a) and (c)), there were more non-migrating cells compared to migrating cells at both days 7 and 14 as expected (**Figure 3 B-C)**. In the monoculture setting where cells of the same type were cultured in both side channels (configurations (b) and (d)), there were also more non-migrating cells compared to migrating cells. Furthermore, there was no significant difference in the number of non-migrating cells between the left and right channels. In co-culture experiments, again there were more non-migrating cells compared to migrating cells. Quantitative analysis confirmed that there were more migrating cells in co-culture settings (configurations (e-g)) as compared to the monoculture settings (configuration (a-d)). In the co-culture settings, the ratio of migrating to non-migrating CMs increased over time.

**Figure 3.**
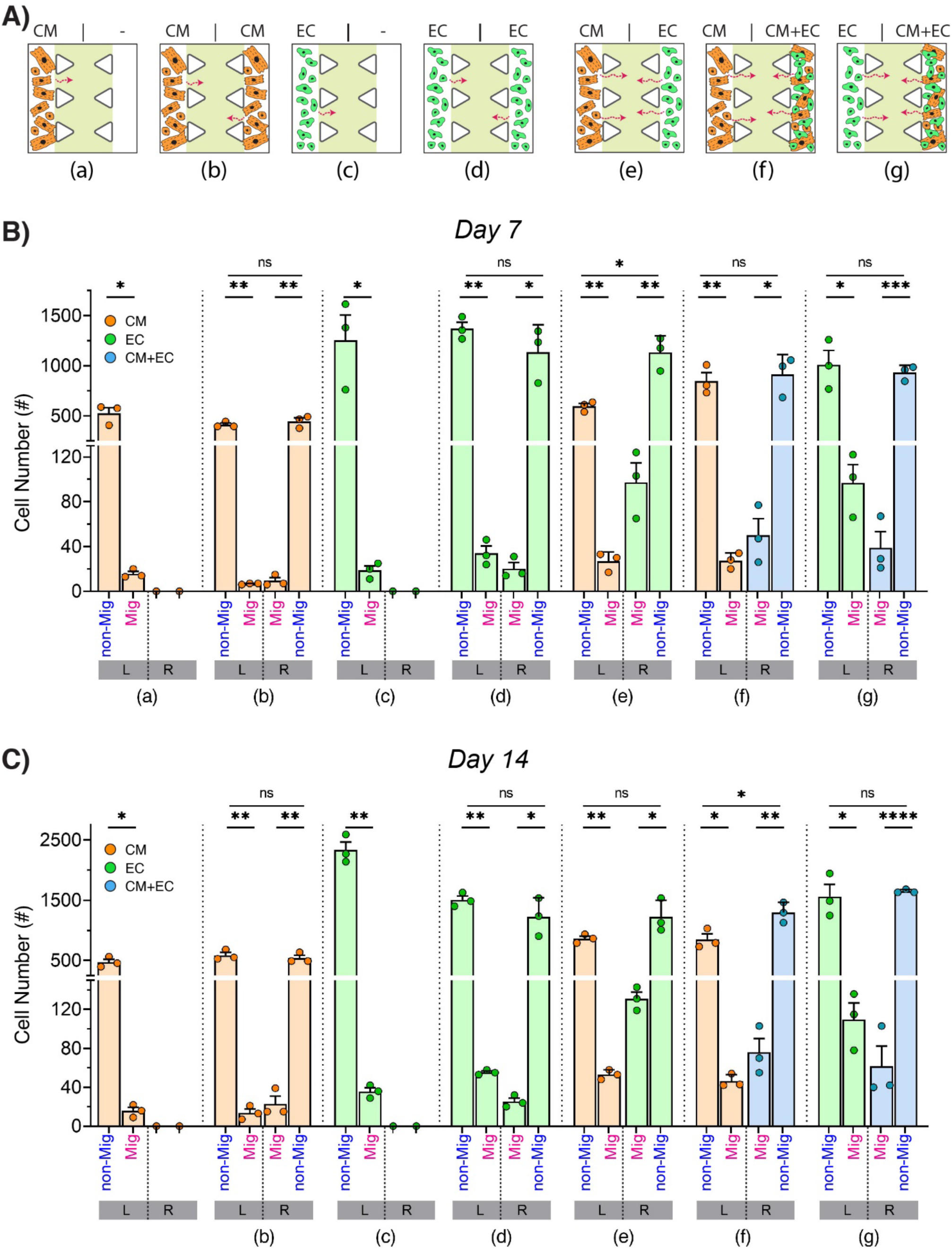
Single cell analysis quantifying the number of migrating cells and the number of non-migrating cells. **A)** Schematic diagram showing the different monoculture and coculture configurations. **B)** Plots showing the number of migrating cells (Mig) vs the number of non-migrating cells (non-Mig) in the left microchannel (L) and the right microchannel (R) at day 7. **C)** Cell migration as in (B) but at day 14. Data represent mean ± SEM and analysed using two-tailed unpaired t-test (*ns* not significant, * *p<0.05*, ** *p<0.01*, *** *p<0.001* and **** *p<0.0001*).

Overall, regardless of cell type, configuration, or mono-vs. co-culture setting, the number of non-migrating cells was always significantly higher compared to the number of migrating cells (**Figure 3 B-C**). This indicated that a subpopulation of cells behaved differently from the bulk population. Usually, it is difficult to identify behavioural differences in individual cells *in vivo* as their properties are masked by the bulk population. Cell heterogeneity plays a vital role in tissue remodelling ^[34,35]^ and an advantage of the current microfluidic device is that it is suited to the visualisation and identification of individual cellular differences. While the majority of neonatal CMs and ECs did not migrate, these data show that a subpopulation possessed properties suited to their migration into a stiffer ECM-like environment and formation of TNT-like structures.

### 2.3. Characterization of neonatal cell sub-populations

We further investigated the behaviour of these CM and EC sub-populations by quantifying their migration characteristics in terms of the number of migrating cells, migration distance and migration speed in different configurations (**Figure 4 A)**. There were no significant differences in the number of migrating CMs or migration distance between different configurations (**Figure 4 B-C**). In the monoculture settings (configuration (a-d)), the number of migrating ECs and the migration distance of ECs were both higher compared to CMs. Taken together, these results indicated that CMs possess lower migration capacity than ECs.

**Figure 4.**
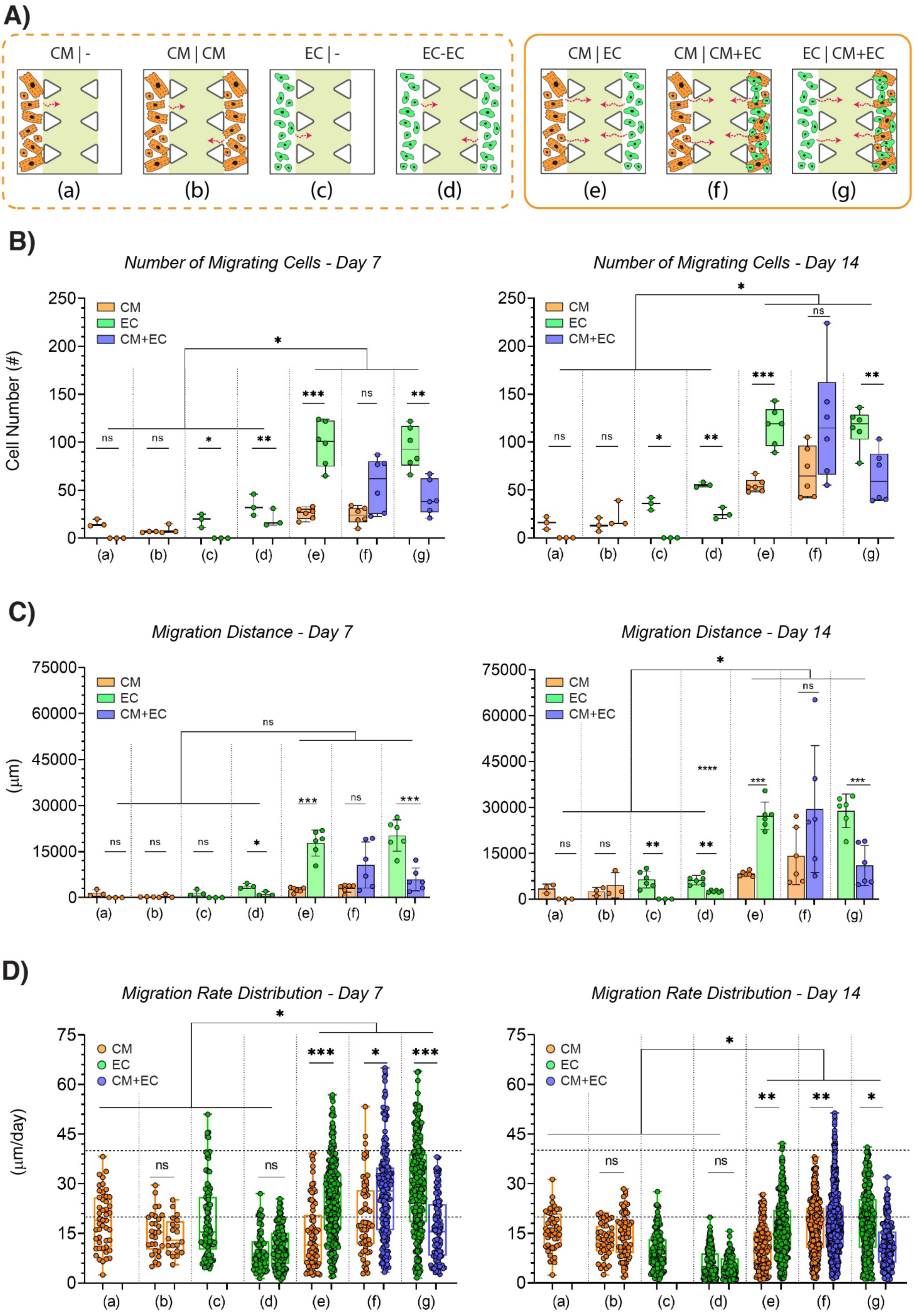
Quantifying migration of neonatal cardiomyocyte and endothelial cells in co-culture microfluidic devices. **A)** Schematic representation of the experimental design. The box with the dashed outline highlights the control experiments (monocultures). The box with the solid outline highlights the coculture experiments. **B)** Box and whisker plots showing the number of migrating cells that moved into the middle channel after 7 and 14 days of culture. **C)** Bar charts showing the migration distance of cells after 7 and 14 days of culture. **D)** Box and whisker plots showing migration rate of cells in which whiskers show the mean and max and middle line represents mean number. Each dot represents migration rate of every single cell. Data represent mean ± SEM and analysed using unpaired two-tailed t-test. (*ns* not significant, * *p<0.05*, ** *p<0.01*, *** *p<0.001* and **** *p<0.0001*).

In general, when cells were grown in co-culture settings (configuration e-g), the number of migrating cells and the migration distance were both significantly higher compared with monoculture settings (configuration (a-d)) regardless of cell type. In configuration (e) and (g) at both days 7 and 14, the number of migrating cells and the migration distance of ECs from an EC-only channel were significantly higher than cells from the opposite channels that contained either CMs alone or a mixture of ECs and CMs **(Figure 4 B-C)**. In configuration (f), at day 7, the numbers of migrating cells from a CM-only channel were lower as compared to cells from the opposite channels that contained a mixture of CM-EC. However, at day 14, no significant difference in the number of migrating CMs were observed between cells from CM-only channel and cells from mixed CM-EC channel. These results indicated that the presence of CMs and ECs in coculture induced the migration of ECs but not CMs. The majority of migration activity was observed in ECs that were not in close proximity to CMs (configuration e-g).

For each randomly selected ROI, the migration rate of individual cells was determined using methods similar to those previously reported ^[36,37]^. In brief, the migration rate was calculated by dividing the migration distance of an individual cell over the duration of the experiment. The migration rates of CMs and ECs were scored as slow (rate<20 μm/day), medium (20<rate<40 μm/day) and fast (rate>40 μm/day). The migration rate for both ECs and CMs in monoculture conditions (configuration (a-d)) were generally slower compared to their co-culture counterparts (configuration (e-g)) (**Figure 4 D**). The majority of CMs at day 7 in CM-EC co-culture configuration (e), migrated at a slow speed, similar to that observed with CMs in monoculture. On the other hand, migration of ECs, was more heterogeneous with rates ranging from slow to fast but the majority at medium speed. Although the migration rates for CMs did not change at day 14, EC migration slowed at day 14. CM migration in configuration (f) was mostly slow at both days 7 and 14 but faster in this configuration than CMs in configuration (e). In configuration (g), where ECs were co-cultured with a CM-EC in the opposite channel, the migration rate for ECs was similar to that in configuration (e). The majority of ECs migrated at medium speed at day 7 and slowed at day 14. Interestingly, the mixed CM-EC cells migrated slower in configuration (g) compared to those in configuration (f), regardless of the culture period. Indeed, the overall migration rate of CM-EC cells from the mixed channel (low to medium speed) was similar to cells in monoculture (low to medium speed).

Microfluidic co-culture systems are excellent tools that enable fine-tuning variables that are not possible with other routine cell-culture systems ^[38]^. They allow users to dissect the impact of these variables at a single cell level. Here, by using a microfluidic co-culture system, we were able to investigate how neonatal CMs and ECs migrate in a systematic and biologically relevant manner. Our results indicated that there was likely a degree of crosstalk between CMs and ECs when they co-exist in culture. Their crosstalk appeared to recruit a sub-population of CMs and ECs to migrate towards each other and form TNT like structures. Sub-populations of migrating cells were found in both CM and EC populations. The subpopulation of CMs exhibited lower migration potential compared to ECs. Furthermore, the crosstalk between CMs and ECs appeared to be stronger at long-range recruitment of ECs that were resistant to migration when in close proximity to CMs.

Crosstalk between CMs and ECs is known to stimulate the secretion of a variety of paracrine signals that regulate a range of cardiac functions such as contractility, metabolism, glucose uptake and cardiac remodelling ^[12,39]^. The crosstalk that led to CM and EC migration that was observed in these experiments were also probably paracrine in nature. It was likely that CMs and ECs secreted soluble factors that diffused throughout the microchip and helped recruit EC and CM sub-populations to migrate towards each other across a distance several times that of a cells diameter.

### 2.4. Spontaneous association of CMs and ECs and formation of organoids on a microchip

Immunofluorescence studies were performed with CM and EC specific antibodies to study possible changes in protein expression induced by coculture on a microchip. At the end of 14 days, cells were fixed and immunolabeled for cardiac α-Actinin (grey) and CD31 (red) (**Figure 5**). CD31, also known as PECAM-1 (platelet/endothelial cell adhesion molecule 1) is an adhesion protein that is highly expressed on ECs ^[40]^. ECs however do not express α-Actinin. CMs express cardiac α-Actinin but not CD31 ^[41]^. As protein expression of these markers on CMs and ECs is well documented ^[40,42]^ and were demonstrated in preliminary studies (**Supporting information**), we used these markers for downstream analysis.

**Figure 5.**
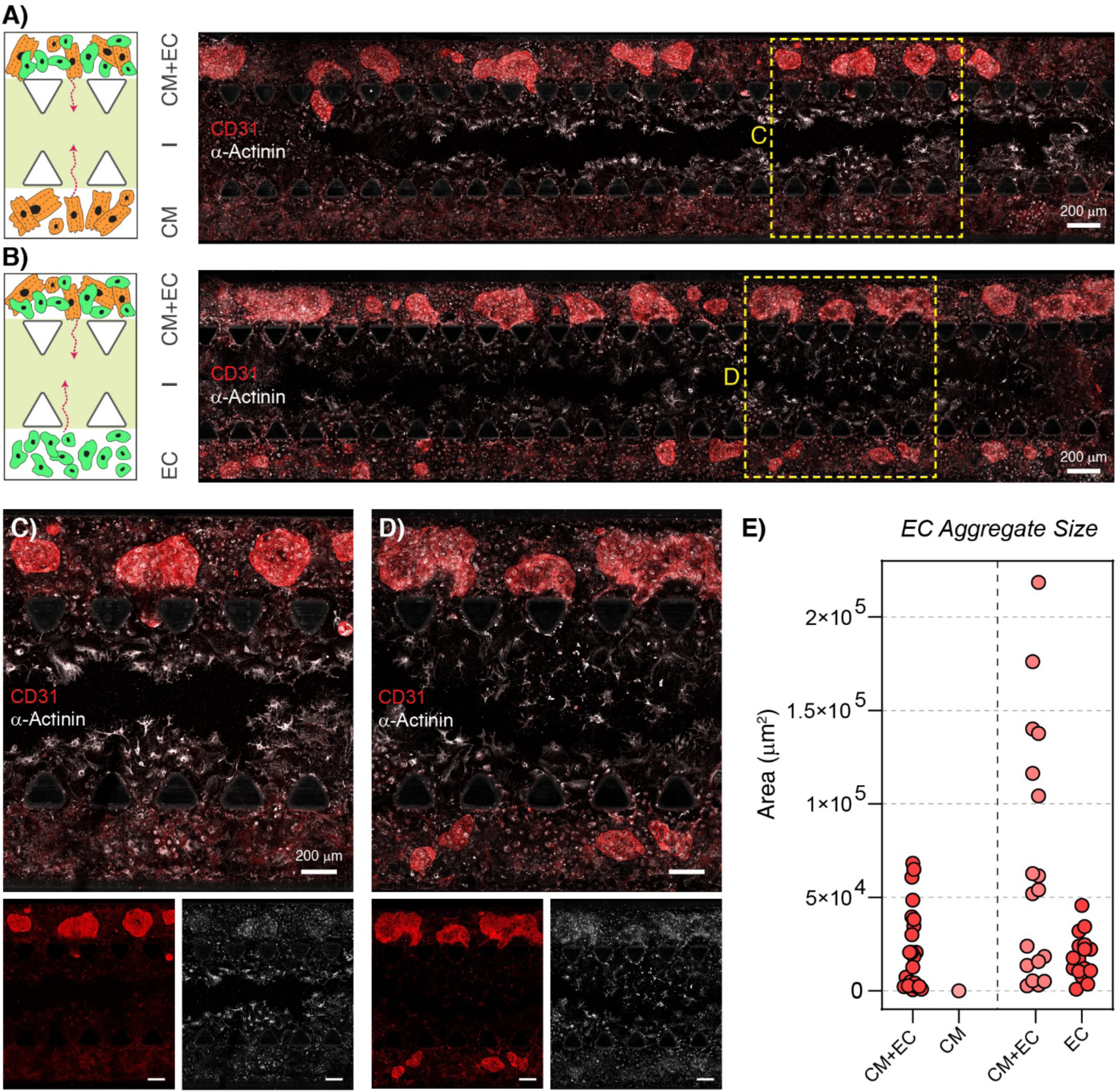
Representative confocal microscopic images demonstrated that non-migrating ECs formed cell aggregates. **A-B)** Tile-scan confocal imaging of microfluidic chips in co-culture conditions. Images shown that cells in side channels that contained ECs have re-organised to form cell aggregates. Endothelial marker CD31 (red) was highly expressed by the cell aggregates. **C-D)** Magnified area boxed in (A-B) showing expression of CD-31 (red) and α-actinin (grey) in cells throughout the microchips. **E)** Plot showing the size of cell aggregates found in Left and Right channels.

As shown in **Figure 5 A-D**, cell aggregates were found in both left- and right-side channels that contained ECs. These cell aggregates were not found in the side channel that contained CMs only (**Figure 5 A**). ECs formed ring-like structures in monoculture conditions when CMs were not present (**Figure S3**). However, in the presence of CMs in the opposite channel, ECs in EC-only channels mostly formed small cell aggregates with few ring-like structures (**Figure 5 B**). This result indicated that formation of EC cell aggregates was likely triggered by soluble factors secreted by CMs diffusing from the opposite side channel across the ECM filled middle channel. Interestingly, some cell aggregates were found to migrate as a cohesive group from a side channel to the middle channel (**Figure 5 C** and **Figure S4**). Migration of cells as a cohesive group, also known as collective cell migration, is a remodelling event that commonly occurs during embryonic morphogenesis and regeneration ^[43,44]^. As shown in **Figure 5 C-D**, cells found throughout the microchip co-expressed α-Actinin and CD31. This indicated that there was a degree of trans-differentiation throughout the co-culture. Some studies have previously reported the trans-differentiation of terminally differentiated endothelial cells into cardiomyocytes in developing mammals ^[28,29,45]^. These results show the plasticity of neonatal CM and EC identity during heart development.

We further measured and plotted the size of cell aggregates, where the size was measured as the cross-sectional area of each cell aggregate (**Figure 5 E)**. The cell aggregates in the mixed CM-EC channels where much larger than EC aggregates in EC-only channel suggestive of CM-EC contact mediated acceleration and growth of cell aggregates.

The major constituents of cell aggregates were ECs (**Figure S4**). In the absence of CMs, ECs formed small endothelial cell aggregates. When co-cultured with CMs, CMs and ECs formed larger mixed cell aggregates. Some of the cells that wrapped around the large cell aggregates exhibited contractile activity typical of beating CMs (**Video S3**). The cell aggregates that were observed here, could be nascent vascular structures ^[46–49]^ that are seen in cardiac organoids ^[50–52]^. Interestingly, no cell aggregate formation was found in the middle channel, indicating that the migrating sub-population of ECs did not participate in the genesis of organoids after migration. The inability of the migrating subpopulation to form cell aggregates was probably due to the low density of cells with cell density likely a critical factor for initiating organoid formation.

CM-EC crosstalk appeared to play a pivotal role in driving ECs to organise into endothelial cell aggregates, both by cell-cell contact and paracrine signalling. Indeed, in our optimization experiment (**Supporting information**), we found that changing half of the culture medium with fresh media every 4 days provided the best culture conditions in terms of CM contractility, probably because the secreted factors from cells could be maintained as long as specific nutrients were not depleted. This result supported the notion that for differentiation studies that require long-term culture, media change should neither be complete or at short intervals (1-2 days) in order to retain a sufficient level of soluble factors produced by cells in culture. The microfluidic device is an ideal setting to investigate the identity and composition of these soluble factors (e.g. cytokines, chemokines, and growth factors).

### 2.5 Cell-cell connections between ECs and CMs

Rearrangement of gap junctions (GJs) between adjacent ECs are a prerequisite for lumen formation ^[53]^. GJs, the main mediator of cell–cell communication, are specific areas at cell boundaries where membranes of adjacent cells contact each other. GJs facilitate the passage of ions and small molecules between adjacent cells ^[54]^. GJs are built by two hemi-channels, called connexons. Each connexon consists of six assembled proteins named connexin (CX). Twenty-one connexin isoforms have been described and the expression of these isoforms is tissue specific. GJs formed between ECs are known to express CX-40, while those in CMs have also been reported to express CX-40^[55,56]^.

A schematic diagram depicts the sequence of events in microfluidic coculture **(Figure 6 A).** Cells migrated from both side channels into the middle ECM channels **(Figure 6 A-i).** Cell aggregates form in the side channels depending on the presence or absence of CM **(Figure 6 A-ii)**, and gap junction molecule expresses at cell boundaries **(Figure 6 A-iii)**. Immunostaining was performed in order to assess if ECs and CMs build and communicate through intercellular connections in addition to paracrine signalling. To this end, ECs and CMs from both mixed-cell channels and single type-cell channels were investigated for GJ marker expression (**Figure 6**). Expression of CX-40 was sparse in channels that contained only CMs (**Figure 6 B**), and in channel that contained only ECs (**Figure 6 C**). Whereas in mixed-cell channels where EC-CM were in close proximity, CX-40 was highly expressed at the interface between CMs and ECs (**Figure 6 D**). These results demonstrated that cell junction connections were formed between ECs and CMs and support the prospect that cell-cell communication occurs between these cell types.

**Figure 6.**
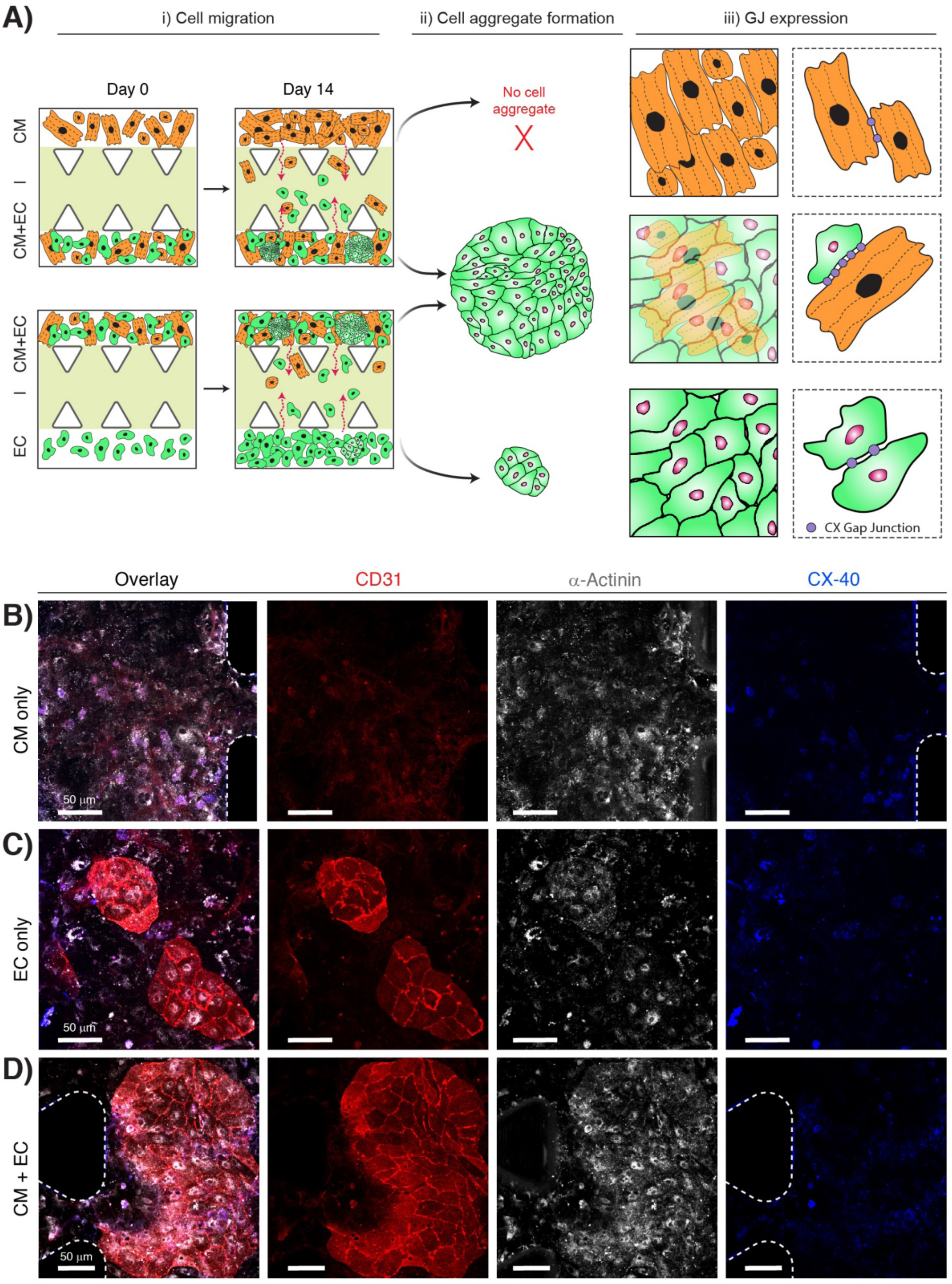
Gap junction formation between CMs, ECs and EC-CMs in co-culture. **A)** Schematic diagram depicting the sequence of events in microfluidic coculture devices: **i)** Cells migrated from both side channels into the middle ECM channels; **ii)** No cell aggregates were found in channel that was seeded with CMs only. Small and large cell aggregates were formed in side channels that were seeded with ECs only or both CMs and ECs, respectively; **iii)** Cartoon representation of gap junction molecule CX-40. **B)** Confocal micrographs showing expression of gap junction molecules CX-40 (blue), CD-31 (red) and α-actinin (grey) in side channel seeded with CMs only. **C)** Confocal micrographs as in (B) but with side channel seeded with ECs only. **D)** Confocal micrographs as in (B) but with side channels seeded with a mixture of CMs and ECs.

The co-cultures were also monitored using microscopy and synchronized CM contractions were observed in the mixed-cell channel with CMs in close contact with ECs (**Video S3**). Electrical coupling between CMs via GJs is a common phenomenon that has been reported as being crucial for synchronized contraction ^[57]^. Our data lends support to this. It is known that endothelial cell heterogeneity exists between different organ systems ^[58,59]^, vessel types ^[22]^, and even between neighboring ECs within the same vessel ^[60]^. This heterogeneity in ECs is partly due to their diverse developmental origins. In this work, we collected all ECs from neonatal hearts. There are reports ^[9,61,62]^ stating that during heart development, ECs in different areas of the heart play specific roles during cardiac development. For example, some ECs from the pericardium migrate into the developing myocardium to participate in vascular-forming events ^[63,64]^. Although speculative, these EC populations might correlate with the migratory ECs that were observed in the microchip. Other ECs (potentially derived from the developing endocardium) do not seem to migrate. These are some of the ECs that become ‘trapped’ as the developing myocardium progresses from being spongy myocardium to compact myocardium ^[9,61]^. Again, although speculative, this population may represent the ECs that formed cell aggregates among non-migrating cells.

EC aggregation and angiogenesis has been shown to precede CM migration in neonatal mouse heart regeneration models ^[65]^. This emphasize that aggregation of ECs guide CMs for migration and organization ^[62]^. If the formation of EC-CM cell aggregates in our device, reflects the ‘aggregation’ process that occurs at the heart compaction stage during neonatal heart development, we may have captured this developmental stage in our microchip. The formation of CM-EC cell-cell connections may facilitate the compaction process and myocardial development ^[66,67]^. It is noteworthy that CM cultures alone did not produce cell aggregates, but these were the preserve of CM: EC co-cultures. We also observed that CMs start contracting early during culture suggesting that the developing myocardium is likely characterised by early contractility (**Video S3**).

## 3. Conclusions

Neonatal myocardium development and maturation are orchestrated by a series of complex cellular interactions that involve cell migration and cell morphogenesis. The sequence and nature of these events require clarity. In this study, a 3D microfluidic co-culture chip was used to study the behaviour of neonatal ECs and CMs to evaluate its potential as a model for studying the development of the vasculature in the neonatal myocardium. First, we observed that from each of the side channels there were subpopulations of cells that migrated toward the ECM mimetic channel, while the majority of cells did not migrate. After 14 days, non-migrating ECs that were not in close contact with CMs formed small cell aggregates. Non-migrating CMs that were not in close contact with ECs, they did not form any cell aggregates. Non-migrating ECs and CMs that were in close contact formed large cell aggregate. GJs were formed at the interface between adjacent cells. Expression of the GJ protein CX-40 was strongest at GJs between CM-EC than EC-EC or CM-CM. Additionally, our microfluidic device enabled us to recapitulate important features of neonatal cardiac development, i.e. collective cell migration, synchronization of CM contraction, and trans-differentiation of ECs and CMs. Taken together, these results demonstrated that the microfluidic coculture system is a useful model, that allows us to recapitulate an early stage of neonatal endothelial-myocardium tissue and offers promise as a means of identifying features and factors that might be useful for future cell therapy to treat coronary artery disease.

## 4. Experimental Section

### 4.1. Neonatal cardiac cell isolation and culture

Mice were housed and bred in the Biological Resource Centre at the Lowy Cancer Research Centre, UNSW Sydney. Neonates were generated by pairing respective strains of male and female mice as shown in the **Figure S1-S3** and **Figure 2–4**. The morning after observing a vaginal plug was designated as day 0.5. All animal experiments were approved by the Animal Ethics Committee of UNSW, Sydney. Hearts were dissected from wild-type C57/BL6 or tdTomato transgenic mice at postnatal day 2 (P2) and were dissociated into single cell suspension using a previously established collagenase enzymatic method ^[8]^. Briefly, the heart tissue was minced into smaller parts and digested using Collagenase II (with a concentration of 263 units/mL) at 37°C. The tissue mixture was gently shaken for 30 min until the hearts were completely dissociated. Cardiac cells were isolated by excluding dead cells using the MACS dead cell removal kit (Miltenyi Biotec, 130-090-101) and were suspended in the optimized culture medium for cardiac cell culture. The medium contained Dulbecco’s Modified Eagle’s Medium (DME)/Ham’s Nutrient Mixture F-12 (Sigma-Aldrich, D6421), and was supplemented with Penicillin/Streptomycin (P/S) (10,000 U/mL) (Thermofisher, Scientific, 10378-016) and 10 % Fetal Calf Serum (FCS) (Scientifix, NZFBS-25). Prior to cell loading, dishes or microfluidic channels were coated with matrigel matrix (Corning, 356237) to enhance attachment and contraction of CM cells. Cardiac cells were cultured in a 5% CO_2_ humidified incubator.

### 4.2. Specific isolation and culture of CMs in a microchip

Primary beating CMs were isolated by MACS (Miltenyi Biotec) according to a reported method ^[68]^. First, primary cardiac cells were isolated from P2 mice using a neonatal heart dissociation kit (Miltenyi Biotech, #130-098-373) according to the manufacturer’s instructions. Next, a neonatal CM Isolation kit (130-100-825, Miltenyi Biotec) was used to sort specific populations of CMs via a gentleMACS machine. CMs were cultured on 10 % matrigel coated microchannels with CM growth medium and maintained in a humidified CO_2_ incubator at 37 °C and 5% CO_2_. DMEM/Ham’s Nutrient Mixture F-12 supplemented with Penicillin/Streptomycin (P/S) (10,000 U/mL) and 10 % FCS was used as CMs medium. The medium was replaced every 4 days. Expression of CM marker cardiac α-Actinin (Sigma-Aldrich, A7811) was assessed by immunolabeling and confocal microscopy (Zeiss LSM 800 confocal microscope). Contraction of CMs was studied using live cell imaging (Zeiss Axio Observer X1 Spinning Disk & TIRF).

### 4.3. Specific isolation and culture of cardiac CD31^+^ EC in a microchip

A neonatal heart dissociation kit (Miltenyi Biotech, #130-098-373) was used to isolate primary cardiac cells from P2 mice hearts. Neonatal cardiac EC isolation kit (Miltenyi Biotec., 130-104-183) was used to further isolate cardiac ECs from the primary cardiac cell population ^[68]^. However, the yield of extraction via this kit was low (~ <10 %). Hence, the isolation protocol was modified. In brief, CD31 microbeads (Miltenyi Biotec., #130-097-418), MS column, and an OctoMACS™ separator were used to extract CD31^+^ cells. ECs were suspended in EGM™-2 Endothelial Cell Growth Medium (Lonza,Basel, Switzerland CC-3162). Cells were cultured on 10 % matrigel coated microchannels and maintained in a humidified CO_2_ incubator at 37 °C and 5% CO_2_. The medium was replaced every 4 days. Expression of EC marker i.e., CD-31 (Abcam, ab28364) and VE-Cadherin (Santa Cruz Biotechnology, sc-9989) were assessed by immunolabeling and confocal microscopy (Zeiss LSM 800 confocal microscope).

### 4.4. Microfluidic device design and fabrication

The microfluidic device was designed according to published methods ^[69–71]^ The device has three channels (10.5 mm in length, and 200 um height), two lateral channels for 2D cell culture, and an inner channel that allows 3D migration of cells (**Figure 1 A-i**). The co-culture microfluidic device was fabricated using a photo lithographic procedure as reported previously ^[72,73]^. Briefly, the photomask was designed via AutoCAD software and then printed on a high-resolution (>8000 dpi) transparency plastic film. Silicon wafers (4 inches, Siegert, DE) were used for mold fabrication. SU-8 photo resist (2100, MicroChem) was patterned on the Si wafer using standard photolithography procedure (Karl Suss MA6 Mask Aligner, SUSS MicroTec, Germany). The patterned wafer was silanized with trichloro (1H, 1H, 2H, 2H-perfluorooctyl) silane (Sigma Aldrich, USA) to render the surface hydrophobic. Subsequently, Polydimethylsiloxane (PDMS) pre-polymer and the curing agent (Sylgard^®^ 184, Dow Corning, USA) were mixed at a 1:10 ratio (curing agent: base), poured into the mould, degassed in a vacuum chamber for 15 min, and baked at 60 °C for 2 h. Finally, PDMS layer was peeled off from the mould, inlets and outlets of 1.5 mm diameter were punched with Harris Uni◻Core (Ted Pella, Inc., USA), and the PDMS layer with microchannel features was bonded onto a coverslip (Thermo Scientific Menzel, 24 × 60 mm) after plasma activation (Harrick Plasma, PDC-002). Microfluidic co-culture devices were washed in Milli-Q water for 20 min followed by a dry autoclave cycle for 20 min.

### 4.5. Co-culture experiment and cell loading onto the chip

Prior to cell loading into the microfluidic device, matrigel solutions (10 % and 40 % in phosphate buffered saline (PBS)) were prepared and were kept within an ice box to prevent gel polymerization prior to device coating. 10 μL of the 40 % matrigel solution was added to the middle channel. The microfluidic device placed in a sterile petri dish was incubated at 37 °C for 30 min. Next, the lateral channels of the microfluidic device were coated with a 10 % matrigel solution and incubated at room temperature for 20 min. In the meantime, primary CMs and ECs were suspended in their corresponding medium and introduced to the micro channels. In the case of CMs + ECs mixture, a mixed culture medium with a ratio of 1:1 was used. Microfluidic chips were kept in a 5% CO_2_ at 37 °C incubator. To prevent the evaporation of media, each microfluidic chip was placed into a humidity chamber made from a petri dish filled with 10 mL of sterile PBS. The culture medium for each cell type was changed every 4 days.

### 4.6. On-chip cell migration analysis and Quantification

Migration of cells from the side channels into the middle channel was studied using Zeiss Axio Observer X1 Spinning Disk & TIRF microscope (Zeiss, Germany) in phase contrast mode via 20X objective. The microscope was equipped with incubation chamber and motorized stage. The incubation chamber was adjusted to operate at 37 °C, 95% humidity, and 5% CO_2_ content. A rectangular array of 18×4 was selected in tile scanning mode and 72 images per chip were taken and stitched together using the ZEN black-ZEISS Efficient Navigation software. In each experiment 3 region of interest (ROI) was selected for quantification analysis (**Figure 1 B**). ImageJ software was used to perform cell tracking, image processing and analysis. The following readouts of cell migration were calculated: i) total number of migrating cell from each side channel into the 3D ECM middle channel, ii) migration distance [μm] (total Euclidean distance from start to end point travelled by every single cell), and iii) distribution of migration rate [μm/day] per cell (calculated by migration distance divided by time). Euclidean distance ^[74]^ or displacement ^[75,76]^ was defined as the linear shortest distance between start and end point of a single cell trajectory (**Figure 1 A-iv**).

### 4.7. Immunofluorescence staining

At the end of a culture period, culture media were removed from the microchannels, and the device was rinsed with PBS for 30 min. The cells were then fixed in 4% paraformaldehyde (PFA) in PBS (w/v) for 15 min at room temperature (RT). Next, cells were permeabilized with Tween-20 for 15 min, washed with PBS and then blocked with 10% donkey serum in PBS for 2 h. Subsequently, cells were incubated in primary antibodies (detailed in **Table 1**) followed by three washes with PBS, and final incubation with secondary antibodies for 1 hr. 4’,6-Diamidino-2-phenylindole, dihydrochloride (DAPI) was used for nuclei staining. Imaging was performed using a Zeiss LSM 800 confocal microscope and the ZEN - ZEISS Efficient Navigation software while the imaging parameters were kept constant during imaging. Images were analysed using FIJI (ImageJ).

**Table 1.** Antibodies used in this chapter.

| REAGENT | SOURCE | IDENTIFIER |
| --- | --- | --- |
| *Primary antibodies* |  |  |
| α-sarcomeric actinin | Sigma-Aldrich | A7811 |
| Anti-CD-31 (Rabbit anti mouse) | Abcam | ab28364 |
| VE-Cadherin | Santa Cruz Biotechnology | sc-9989 |
| Connexin 40 Polyclonal | Thermo Fisher Scientific | 36-5000 |
| Anti-Connexin-43 | BD Biosciences | 610062 |
| DAPI | Invitrogen | H3569 |
| *Secondary antibodies* |  |  |
| Donkey anti Mouse-488 | Invitrogen | A21202 |
| Donkey anti Rat-488 | Thermo Fisher Scientific | A21208 |
| Donkey anti Rat-650 | Invitrogen | A10043 |
| Chicken anti Mouse-647 | Invitrogen | A21463 |
| Rabbit anti Mouse-555 | Invitrogen | A21427 |
| Donkey anti Rabbit-555 | Thermo Fisher Scientific | A31572 |

### 4.8. Statistical analysis

Statistics were performed using Graph Pad Prism software (Version 8.0.2, Inc., La Jolla, CA, USA). Sample size is indicated within corresponding figure legends and all data are presented as mean ± SEM. p-values were determined using unpaired two-tailed student’s t-test. In all figures, ns represents not significant, * represents p < 0.05, ** represents p < 0.01, *** represents p < 0.001, and **** represents p < 0.0001.

## Author’s Contribution

H.T, V.C and P.C designed the study. H.T performed the experiments including microfluidic device fabrication, heart tissue dissection, cardiac cell isolation, and microscopic analysis. Y.C.K and P.R performed animal mating. J.E.P advised on refining experiment design. H.T analysed and interpreted the data. H.T, P.C and V.C wrote the manuscript and J.E.P revised the manuscript. P.C and V.C supervised this study. J.E.P, V.C and P.C acquired funding.

## Acknowledgements

H.T. thanks the financial support from Swinburne University of Technology through SUPRA scholarship, and the Stem Cell Group at Adult Cancer Program, UNSW Sydney. Funding for this research was partly provided through an Australia Research Council Discovery Project Grant DP170103704 and UNSW medicine. J.E.P and V.C acknowledge funding from the National Health and Medical Research Council of Australia (APP1061593 and APP1139811).

## Conflict of Interest

There are no conflicts to declare.

## Notes

### Competing Interest Statement

The authors have declared no competing interest.

