## Supplemental File for "Capturing Neonatal Cardiomyocyte – Endothelial interactions on-a-Chip"

### Contents

**Overview of experiments design and optimization for co-culture on-a-chip**

Isolation of cardiac cells and optimization of culture medium

Viability of isolated cardiac population

Isolation of CMs from cardiac population and proliferation assessment

Isolation and purification of ECs from cardiac population

**Supplementary Figures**

**Figure S1.** Experimental procedure for preparation and isolation of cardiac cells and culture into microchip**.**

**Figure S2.** Experimental procedure for preparation and isolation of CMs and culture into microchip.

**Figure S3.** Experimental procedure for preparation and isolation of ECs and culture into microchip.

**Figure S4.** Representative confocal images showing a cell aggregate undertaken collective cell migration.

**Supplementary Movies**

**Video S1.** Contraction of neonatal cardiac cells after 7 days.

**Video S2.** Contraction of neonatal CMs after 7 days.

**Video S3.** Contractile activities of cell aggregates after 7 days.

#### Overview of experiments design and optimization for co-culture on-a-chip

##### Isolation of cardiac cells and optimization of culture medium

Wild type C57/BL6 mice were time mated to generate wild-type neonates (**Figure S1 i**). Harvested neonates were digested using a collagenase and cultured on Matrigel coated plates. Commercially available Miltenyi kit was also employed for neonatal cardiac cell isolation and results compared. Microscopic observation showed that there were more areas of contracting cells in the cultures isolated using the Miltenyi kit. Hence, Miltenyi kits were used for all experiments. Next, due to variability in reports regarding culture media, three culture media (α-MEM, DMEM, and DMEM/Ham's F-12) were tested. In the media containing DMEM/Ham's F-12, cardiac cells reached confluency within the shortest timeframe while their contractions were more synchronized. Finally, cardiac cells were cultured on coated plates at different seeding densities ($10\times10^5$, $5\times10^5$, $2.5\times10^5$, $1\times10^5$, $0.5\times10^5$ and $0.25\times10^5$ cells/ml) in order to find the minimum number of beating cells needed for culture in the microfluidic chip. The cells were monitored every 12 hrs for 7 days. Cardiac cells at a seeding density of $0.5\times10^5$ cells/ml started beating after 36 hours of culture. These optimized conditions (isolation method, cell number, and culture media) were used for culture on the microfluidic chip.

##### Viability of isolated cardiac population

The whole heart contains a heterogeneous population of cells including CM, EC, fibroblast etc. (**Figure S1 ii**). To assess if the whole population of cardiac cells were still viable after isolation, and to assess if they can be cultured on the microfluidic chip, these cells were cultured in the side channels of the microfluidic chip and allowed to migrate into the middle channel (**Figure S1 iii**). The cell culture on-chip was imaged at day 1 (**Figure S1 iv**) and day 7 (**Figure S1 v**). It was observed that after a week, cells proliferated and covered the channel as a confluent cell layer (**Figure S1 v**). In some areas, beating CMs were observed (**Video S1**). After 2 weeks, cells were fixed and labelled with both cardiac and endothelial biomarkers. **Figure S1 (vi-vii)** shows the morphology of CD31 expressing EC, α-Actinin expressing CMs, and the nuclear stain DAPI. Imaged areas show cardiac cell migration from the side channels to the middle channel.

**Isolation of CMs from cardiac population and proliferation assessment**

Neonatal CM cells from wild type mice were isolated (**Figure S2 i**) using MACS (**Figure S2 ii**). Next, CMs were loaded into the microfluidic channels coated with a layer of 10% matrigel (**Figure S2 iii**). 24 hours after cell loading, the majority of CMs showed contractile properties and the contraction pattern became synchronized after 48 hrs in culture. Phase contrast microscopy images showed that the number of CMs increased after 7 days of culture indicating that these CMs had proliferated (**Figure S2 iv-v**). After immunolabeling, CMs were found to express typical cardiomyocyte biomarkers (**Figure S2 vi**), in particular, sarcomeric α-Actinin at day 14 (**Figure S2 vii**). Contraction of neonatal CMs was observed after day 7 (**Video S2**).

**Isolation and purification of cardiac ECs**

Finally, cardiac endothelial cells were separated using magnetic beads (**Figure S3 i-iii**). These cells express CD31. Purity of isolated ECs was assessed by FACS and noted to be 86% (**Figure S3 iv**). Representative images show cells that were immunolabeled by CD31. Confocal imaging confirmed the expression of CD31 (**Figure S3 v**).

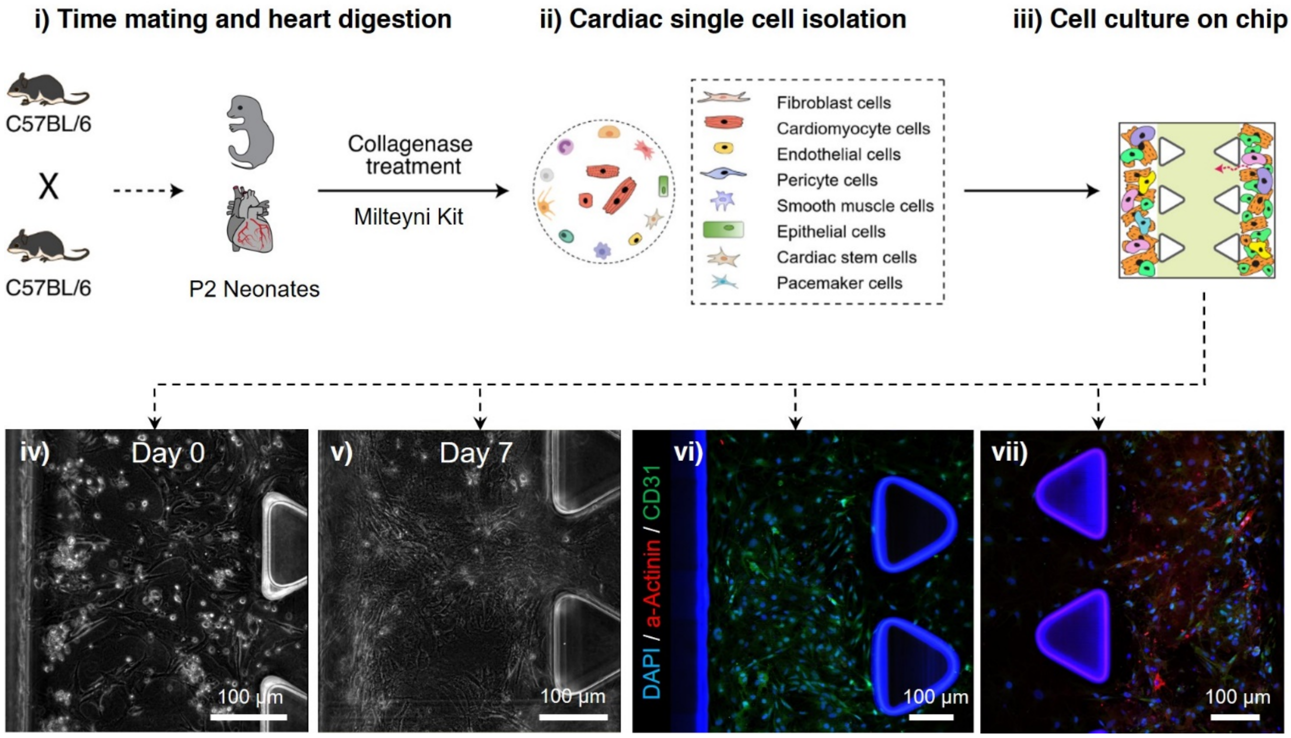


**Figure S1. Experimental procedure for preparation and isolation of cardiac cells and culture on the microchip.** i) Neonatal (P2) heart tissues were digested using collagenase, ii) Primary Neonatal (P2) cardiac cells were isolated, and iii) the entire cardiac cell population was loaded into the microfluidic chip. iv-v) viability, proliferation and contraction of cells was monitored using live cell imaging after 7 days. vi-vii) Immunolabeling of endothelial and cardiomyocyte markers show the expression of CD31 in green and cardiac α-Actinin in red, respectively. Nuclei were stained with DAPI (blue). Imaged areas show cardiac cells in the side channels migrating into the middle channel.

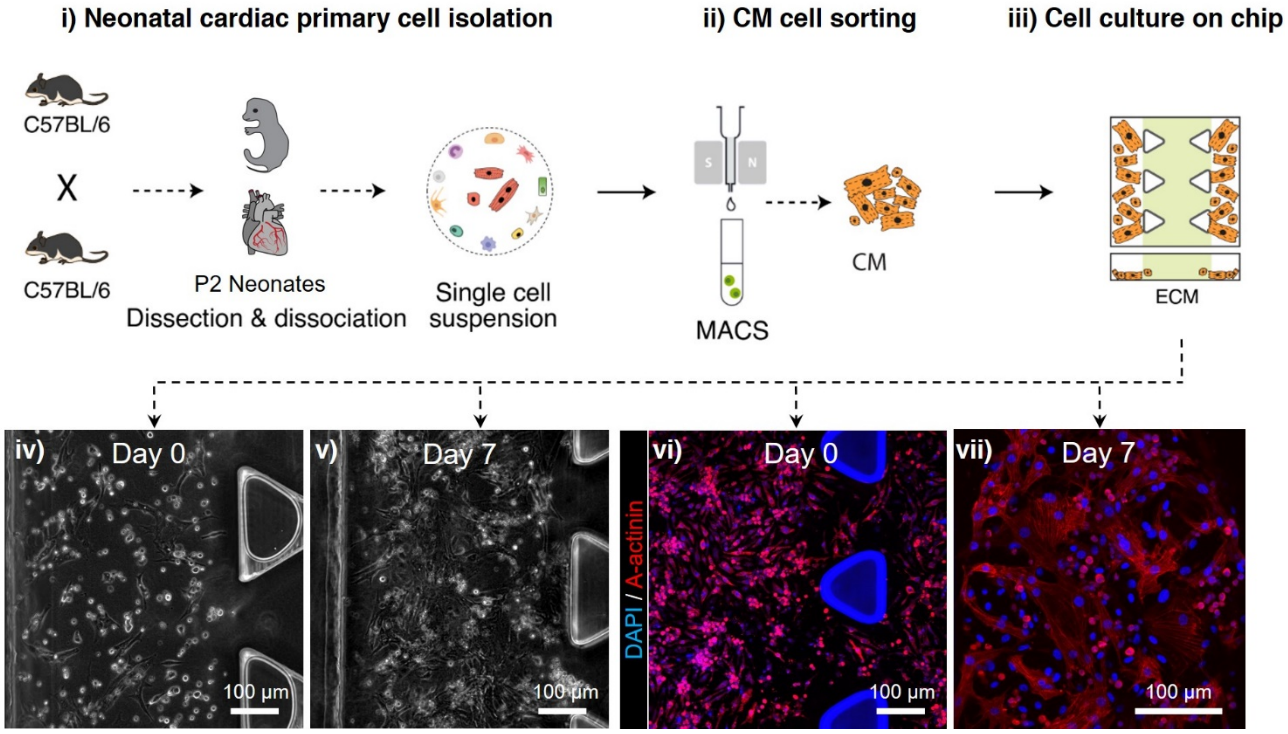


**Figure S2. Experimental procedure for preparation and isolation of CMs and culture on the microchip.** i) Neonatal (P2) heart tissues were digested using a Miltenyi kit. ii) Magnetic-activated cell sorting (MACS) was used to isolate primary neonatal CM cells; and iii) cells were loaded into the microfluidic chip. iv-v) viability, proliferation and contraction of CMs were observed using live cell imaging at day 0 and day 7. Images were taken from the side channel. vi-vii) CMs were immunolabelled for cardiac α-actinin expression (in red). The nuclei were stained with DAPI (blue).

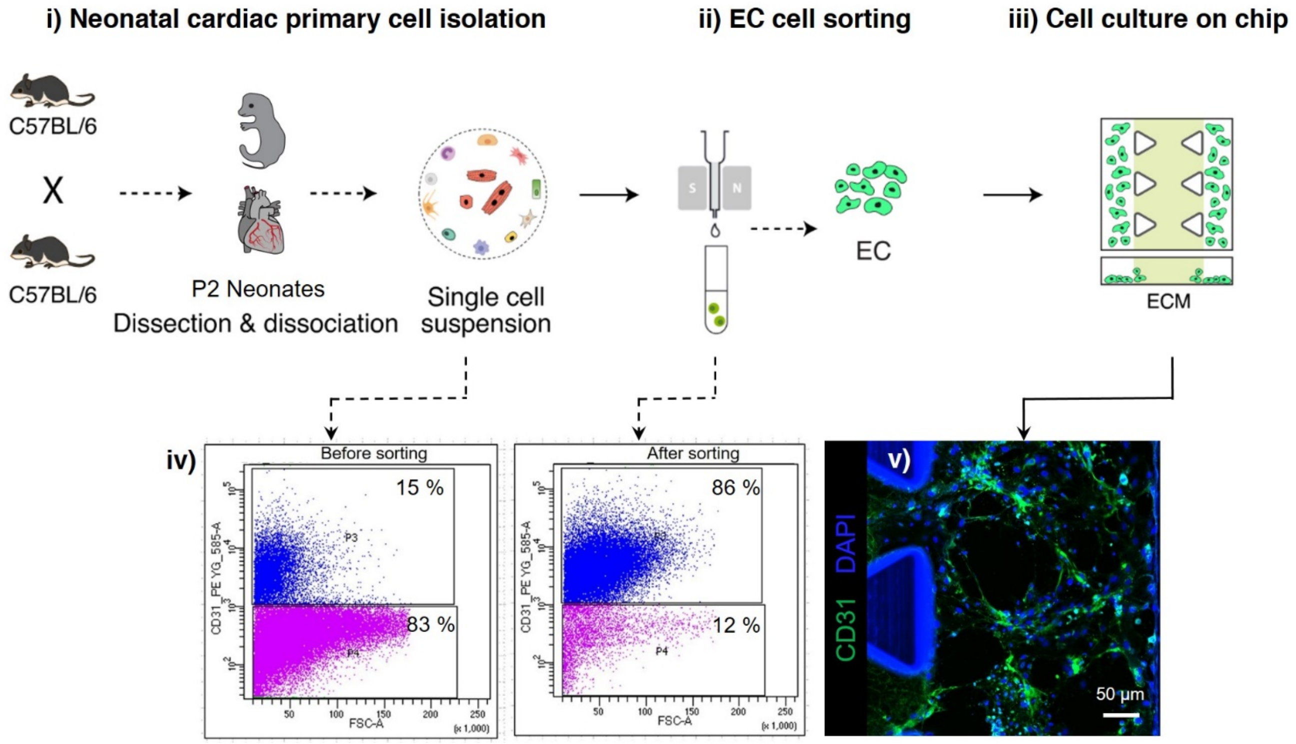


**Figure S3. Experimental procedure for preparation and isolation of ECs and culture on the microchip.** i) Neonatal (P2) heart tissues were digested using a Miltenyi kit. ii) MACS was used to isolate primary neonatal endothelial cells; and iii) cells were seeded into the microfluidic chip. iv) fluorescence activated cell sorting (FACS) analysis method was used to assess the isolation purity: pre-sort purity (CD31+) was ~15 % and post-sort purity ~86 %. v-iv) Immunolabeled ECs expressed CD31 (green). Cell nuclei were stained with DAPI (blue). Images were taken from the side channel.

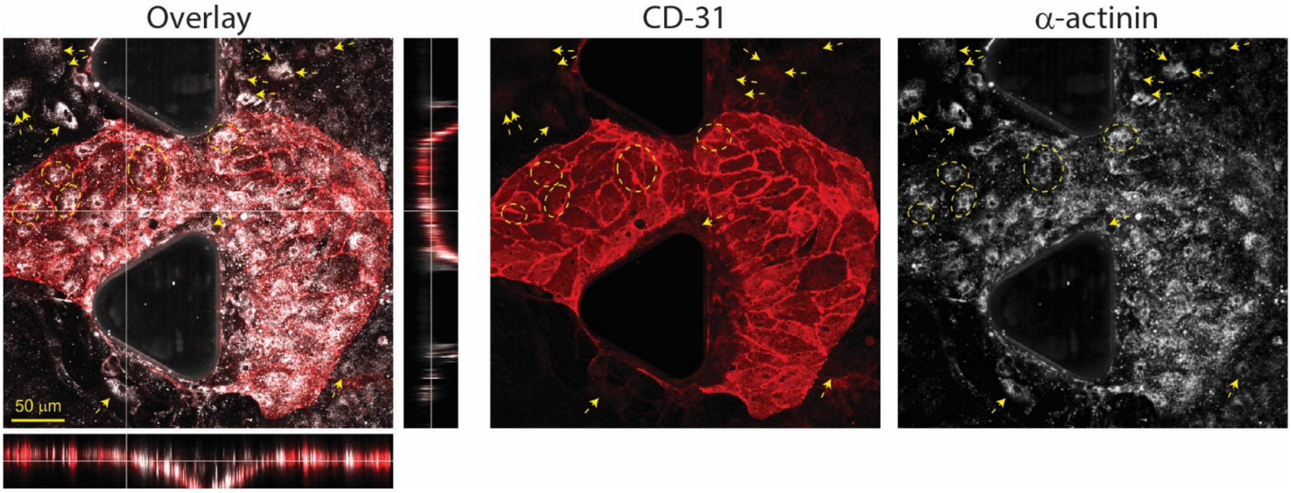


**Figure S4. Representative confocal images showing a migrating cell aggregate.** The cell aggregate was formed in a side channel that housed both CMs and ECs. The aggregate was observed to migrate towards the ECM channel as a whole. Overlay image shows a z-scan confocal image of the cell aggregate. The cell aggregate was stained for CD31 (red) and α-Actinin (grey) expression. Yellow dotted lines highlight comparable regions in individual channels. Aggregates are formed from both CD31+ and α-Actinin+ cells.

**Video S1.** Contraction of neonatal cardiac cells after 7 days.

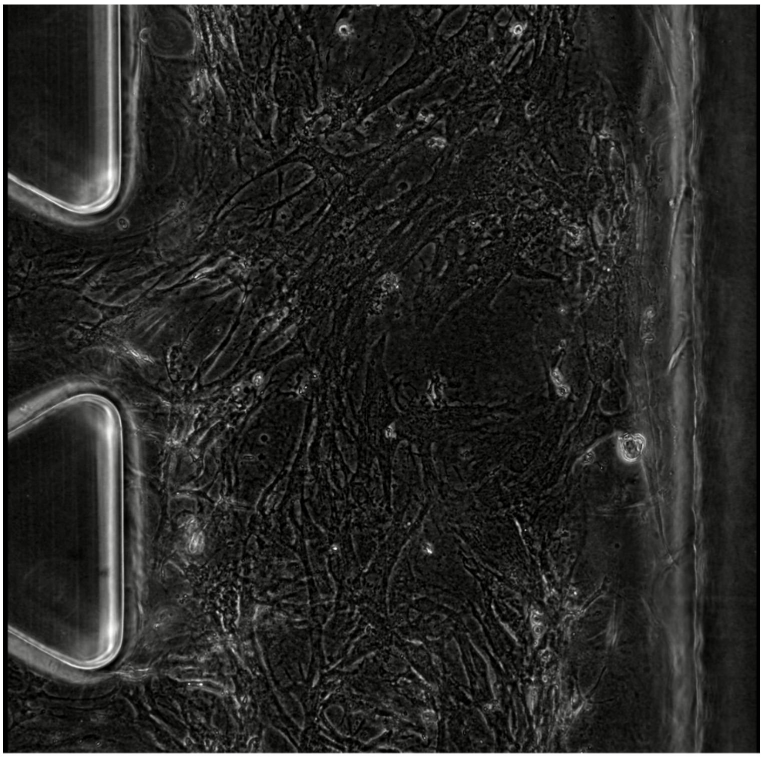

**Video S2.** Contraction of neonatal CMs after 7 days.

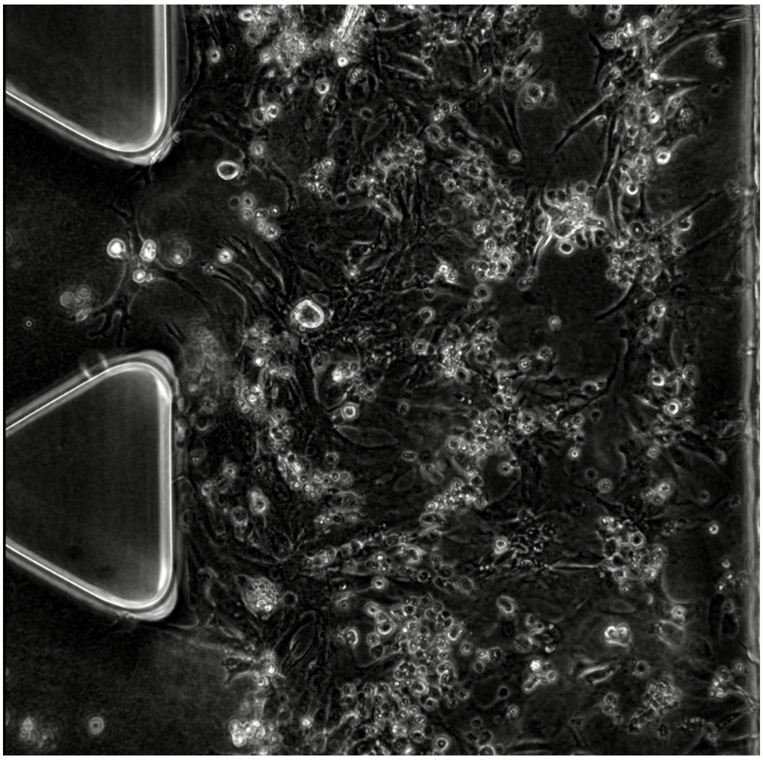

**Video S3.** Contractile activities of cell aggregates after 7 days.

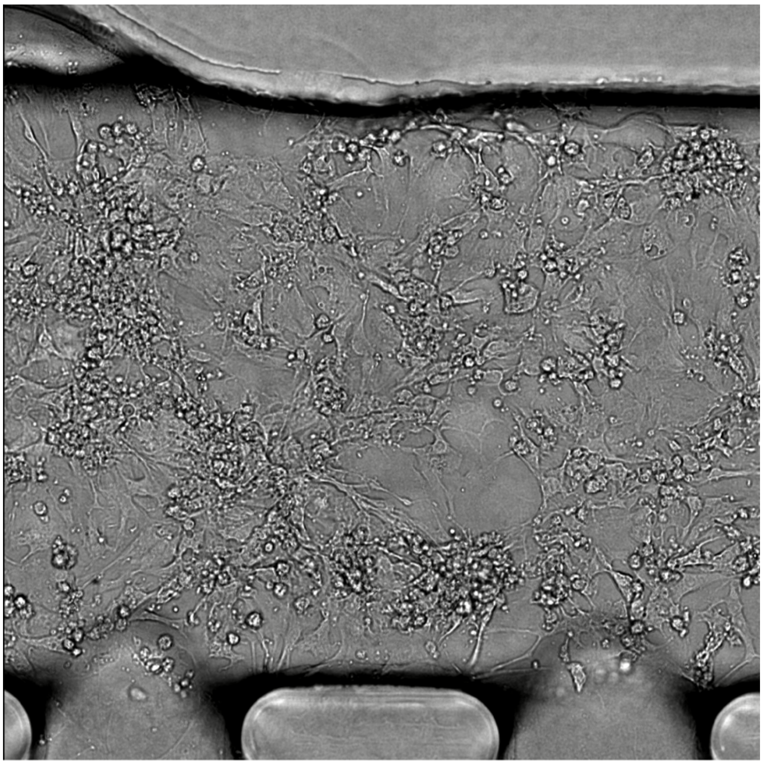
